# Knockout of AMD-associated gene *POLDIP2* reduces mitochondrial superoxide in human retinal pigment epithelial cells

**DOI:** 10.1101/2022.07.05.498910

**Authors:** Tu Nguyen, Daniel Urrutia-Cabrera, Luozixian Wang, Jarmon G. Lees, Jiang-Hui Wang, Sandy S.C. Hung, Alex W. Hewitt, Thomas L Edwards, Shiang Y. Lim, Chi Luu, Robyn Guymer, Raymond C.B. Wong

## Abstract

Genetic and epidemiologic studies have significantly advanced our understanding of the genetic factors contributing to age-related macular degeneration (AMD). In particular, recent expression quantitative trait loci (eQTL) studies have highlighted *POLDIP2* as a significant gene that confers risk of developing AMD. However, the role of *POLDIP2* in retinal cells such as retinal pigment epithelium (RPE) and how it contributes to AMD pathology are unknown. Here we report the generation of a stable human RPE cell line with *POLDIP2* knockout using CRISPR/Cas, providing an *in vitro* model to investigate the functions of *POLDIP2*. We conducted functional studies on the *POLDIP2* knockout cell line and showed that they retained normal levels of cell proliferation, cell viability, phagocytosis and autophagy. Also, we performed RNA sequencing to profile the transcriptome of *POLDIP2* knockout cells. Our results highlighted significant changes in genes involved in immune response, complement activation, oxidative damage and vascular development. We showed that loss of *POLDIP2* causes a reduction in mitochondrial superoxide levels, which is consistent with the upregulation of the mitochondrial superoxide dismutase *SOD2*. In conclusion, this study demonstrates a novel link between *POLDIP2* and *SOD2*, which supports a potential role of *POLDIP2* in regulating oxidative stress in AMD pathology.

## Introduction

Age-related macular degeneration (AMD), the most common cause of irreversible vision loss amongst people over 50 years old in developed countries, ^1,2^ is characterised by RPE degeneration and photoreceptor cell death. The RPE is critical to retinal homeostasis and some important roles of RPE include phagocytosis of photoreceptor outer segments, scavenging for damaged reactive oxygen species (ROS), and delivery of blood-derived nutrients to photoreceptors ^3,4^. Emerging evidence in recent years suggests that oxidative stress-induced mitochondrial damage in the RPE contributes to development of AMD ^5–7^.

Genome-wide association studies (GWAS) have contributed to our understanding of genetic associations to AMD and identified >60 single nucleotide polymorphisms (SNPs) implicated in AMD ^8–10^, including *CFH* and *ARMS2/HTRA1* loci that confer major susceptibility ^11–13^. In addition, other genomic methods have been developed to complement GWAS in identifying and confirming variants associated with diseases, such as expression quantitative trait loci (eQTL), transcriptome-wide association study (TWAS), and eCAVIAR ^10,14–16^. These techniques integrate genomic and transcriptomic data sets to confirm whether causal genes found in GWAS studies are driving disease association (reviewed in ^17^). In particular, using eQTL and TWAS, a recent study highlighted *POLDIP2* at the *TMEM97/VTN* loci as a significant target gene associated with AMD ^14^. However, the function of *POLDIP2* in the retina remains poorly understood. Our previous study has shown that *POLDIP2* is highly expressed in human RPE/choroid ^28^. Unravelling the biological roles of *POLDIP2* in RPE is critical to advancing our understanding of AMD pathogenesis.

*POLDIP2* encodes a multifunctional protein that localises in both the nucleus and the mitochondria ^18^. Several studies have identified *POLDIP2* as a significant gene for AMD susceptibility ^10,14,16^ and it has been associated with vascular and neurodegenerative diseases ^19,20^. *POLDIP2* has been reported to play a role in a wide range of physiological and cellular processes ^18^. Previous mouse studies showed that homozygous *Poldip2* knockout was embryonic lethal, *Poldip2*-/-embryos were significantly smaller than WT, and mouse embryonic fibroblasts (MEFs) exhibited reduced growth ^21^. In addition, heterozygous knockout mice displayed lower levels of H_2_O_2_ production, which increased aortic extracellular matrix, increased vascular stiffness and impaired contractility, thus demonstrating that *Poldip2* expression is necessary for vascular structure and function ^22^. Poldip2 has also been shown to be an oxygen-sensitive protein and regulates cell metabolism and mitochondrial function ^23^. Poldip2 expression is downregulated by hypoxia and in cancer cells, leading to repression of lipoylation of the pyruvate and α-ketoglutarate dehydrogenase complexes and mitochondrial dysfunction. Interestingly, *POLDIP2* dysfunction has been implicated in Alzheimer’s disease, including metabolic and oxidative stress, neuroinflammation, as well as abnormal microvasculature and extracellular deposits ^24,25^. Overexpression of *POLDIP2* resulted in defective autophagy leading to increased Tau aggregation, whereas *POLDIP2* downregulation decreased ROS-induced Tau aggregation ^20^. However, there is no known study on the function of *POLDIP2* in the retina and its role in the development of AMD.

Recent advances in CRISPR technology offer exciting opportunities to manipulate genes and accelerate functional study of AMD-associated genes. The combined use of Cas9 endonuclease and a single-stranded guide RNA (sgRNA) can target and cleave specific DNA sequences, thereby creating a double-stranded break and deletions (indels) to knockout genes ^26^. Alternatively, a catalytically inactive Cas9 (dCas9) can be coupled with a transcriptional repressor domain, such as Krupper-associate box (KRAB), to repress the expression of a target gene, termed CRISPR interference (CRISPRi) ^27^. Together, these CRISPR/Cas9 systems provide useful tools to perform loss-of-function study for AMD-associated genes in the RPE.

Using CRISPR/Cas9, here we report the generation of a human RPE cell line ARPE-19 with *POLDIP2* knockout. We showed that *POLDIP2* knockout resulted in upregulation in *SOD2* levels and decreased levels of mitochondrial superoxide. Also, our results highlighted the effect of *POLDIP2* loss on the transcriptome profile of RPE, and discovered upregulation of genetic signals related to immune response, oxidative damage and vascular development.

## Results

### Evaluation of CRISPRi and CRISPR knockout of *POLDIP2* in ARPE-19

We first tested the use of CRISPRi to knockdown *POLDIP2* in ARPE-19, using an ARPE-19 cell line overexpressing dCas9-KRAB we reported previously (ARPE-19-KRAB) ^28^. To induce knockdown of *POLDIP2* expression, we designed 2 sgRNAs that target the proximity of the transcription start site (TSS) of the *POLDIP2* gene (Supplementary figure 1). We transfected 2 different concentrations of sgRNAs, 360ng and 1000ng, into ARPE-19-KRAB. The efficiency of the 2 sgRNAs in knocking down *POLDIP2* expression was assessed using RT-qPCR. The results showed that sgRNA1 could not knockdown *POLDIP2* (360ng: 1.15±0.12 compared to mock; 1000ng: 1.15 compared to mock, Supplementary figure 2), whereas sgRNA2 could repress *POLDIP2* expression level by ~24% (360ng: 0.77±0.05 compared to mock; 1000ng: 0.76±0.15 compared to mock, Supplementary figure 2). Since the CRISPRi-mediated knockdown level observed was mild and likely insufficient for functional studies, next we focused on the use of CRISPR/Cas9 to knockout *POLDIP2* in ARPE-19.

To induce *POLDIP2* knockout, we transduced ARPE-19 with lentiviruses carrying sgRNA targeting the coding sequence of *POLDIP2*). Following antibiotic selection, we generated a stable ARPE-19 cell line with *POLDIP2* knockout (POLDIP2 KO). POLDIP2 KO cells retained similar morphology to ARPE-19 wild type (WT) (Supplementary figure 4). Critically, Sanger sequencing confirmed the presence of indels at the target site in *POLDIP2* CDS (Figure 1a), with a 70% indel percentage in POLDIP2 KO compared to WT (Figure 1b). We analysed the levels of *POLDIP2* gene expression using RT-qPCR and found a ~80% reduction of *POLDIP2* levels in POLDIP2 KO (0.22±0.01 compared to WT, Figure 1c). Also, western blot analysis showed an absence of POLDIP2 protein expression in POLDIP2 KO samples (Figure 1d). Finally, we performed a short tandem repeat (STR) analysis and confirmed that the knockout cell line originated from the parental ARPE-19 cell line (Supplementary figure 3) ^28^. Collectively, these results showed that we have successfully generated an ARPE-19 cell line with *POLDIP2* knockout.

**Figure 1.**
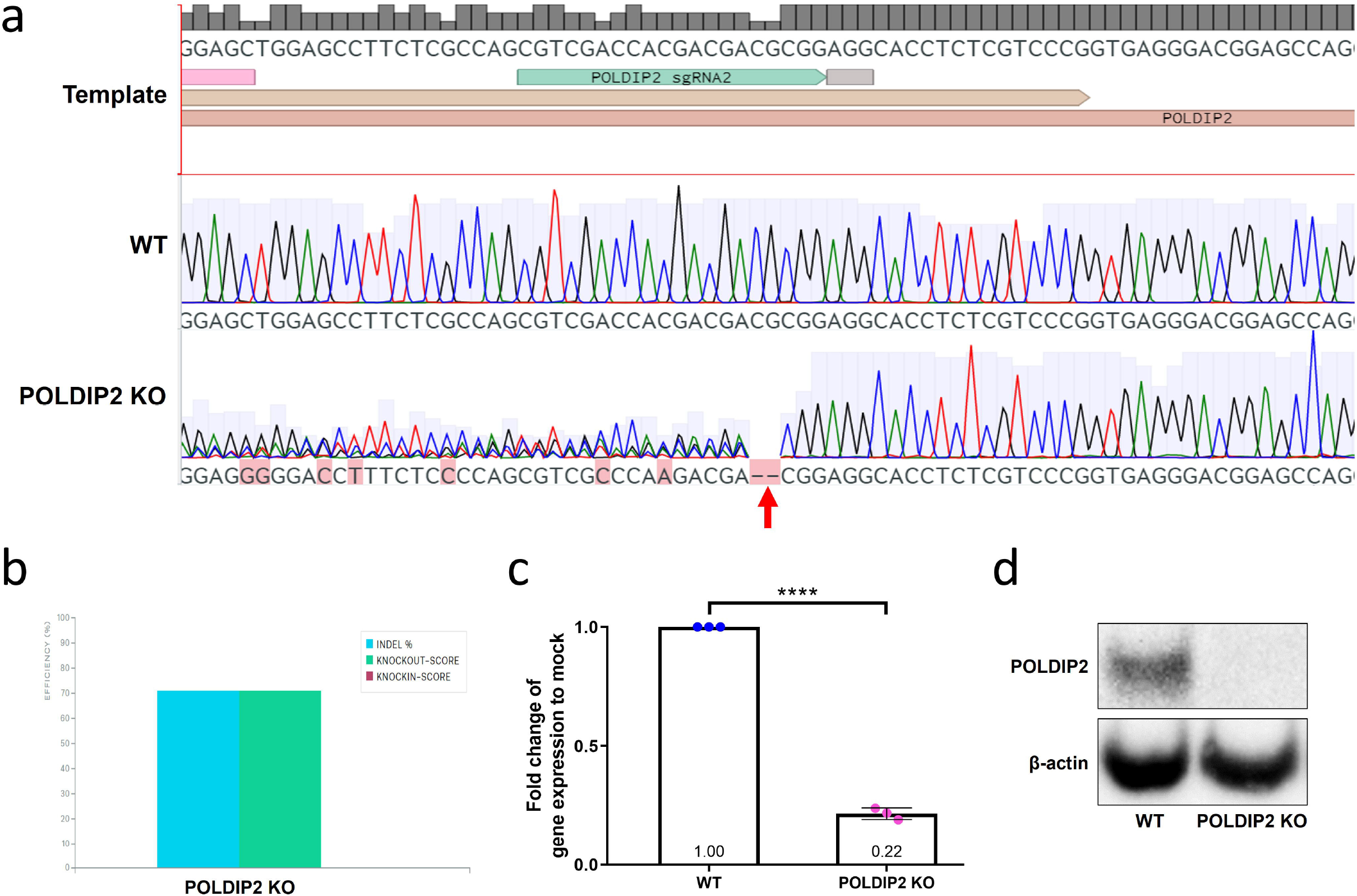
Characterization of POLDIP2 KO cell line. a) Sanger sequencing shows small indels in the coding sequence of *POLDIP2* in the POLDIP2 KO cell line, as indicated by the red arrow. b) Quantification of indel percentage in knockout cell line compared to WT. c) RT-qPCR analysis of *POLDIP2* repression using CRISPR KO. Values expressed as mean ± SEM, n=3. **** p<0.0001. d) Western blot analysis of POLDIP2 protein repression.

### Functional studies of *POLDIP2* on ARPE-19 cells

Phagocytosis is an important function of RPE to degrade reactive oxygen species (ROS) and maintain retinal homeostasis. Using POLDIP2 KO, we assessed the effect of *POLDIP2* on RPE phagocytosis. WT and POLDIP2 KO cells were incubated with FITC fluospheres and their phagocytosis ability was analysed by quantification of FITC+ cells using flow cytometry. Our results showed that the POLDIP2 KO cell line retained the ability to phagocytose FluoSpheres (Figure 2a). The proportion of FITC+ in POLDIP2 KO cells was 34.53±3.19%, compared to 34.27±1.12% in WT, which suggested the levels of phagocytosis between WT and POLDIP2 KO cells were similar (Figure 2b). Our results demonstrated that *POLDIP2* knockout did not affect the phagocytic ability of ARPE-19 cells.

**Figure 2.**
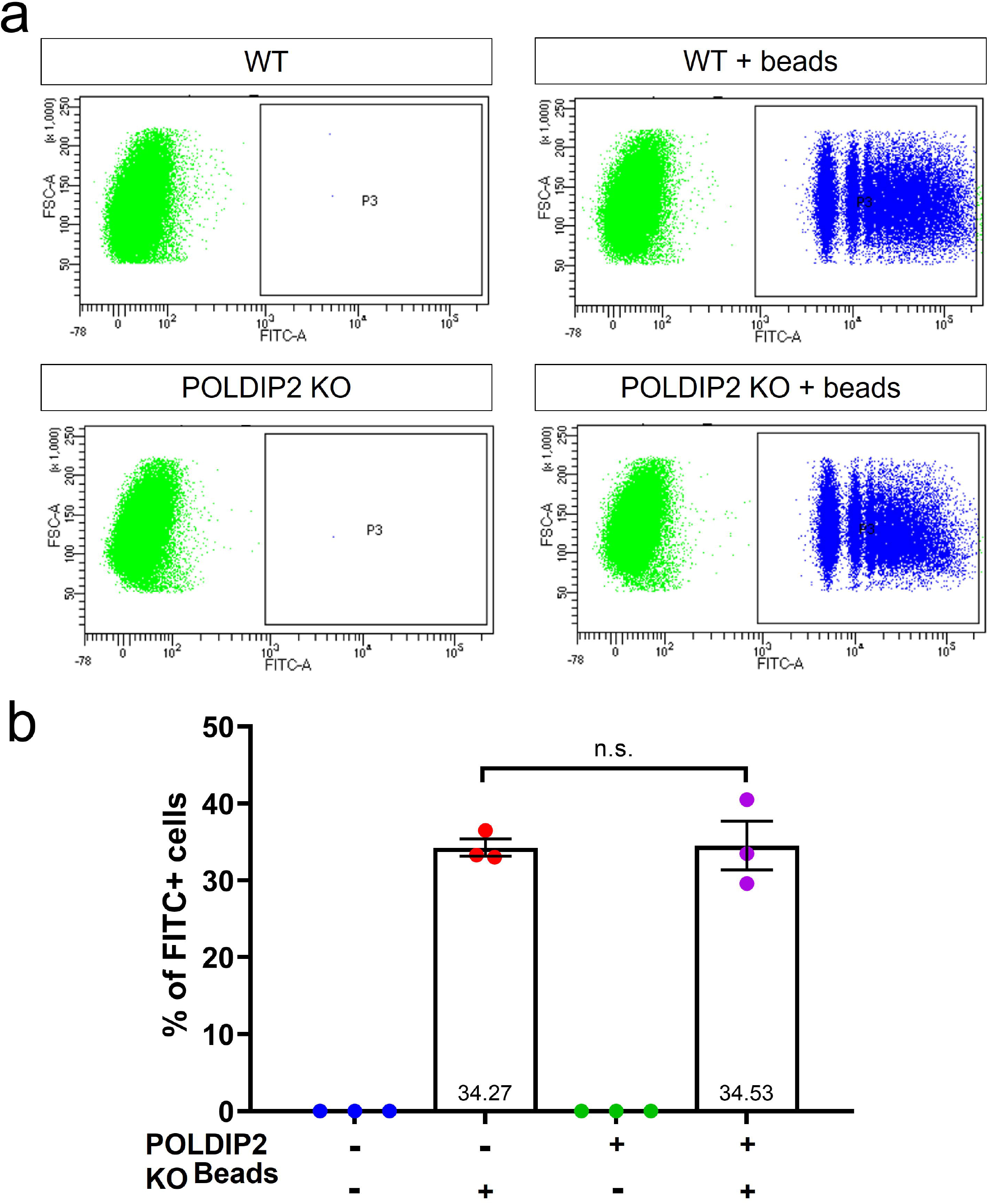
POLDIP2 KO cells show normal levels of phagocytosis. a) Flow cytometry analysis of phagocytosis in WT and POLDIP2 KO treated with or without FITC+ fluospheres. b) Pooled quantification results of n=3 biological repeats. Error bars represent SEM. n.s. not significant.

Next, we investigated whether loss of *POLDIP2* would affect proliferation of ARPE-19. We showed that the POLDIP2 KO cell line showed a comparable growth rate to WT (POLDIP2 KO: R^2^=0.87; slope=3.47; WT: R^2^=0.87; slope=2.89, Figure 3a). Overall, our results showed that the effect of *POLDIP2* loss on cell proliferation is negligible.

**Figure 3.**
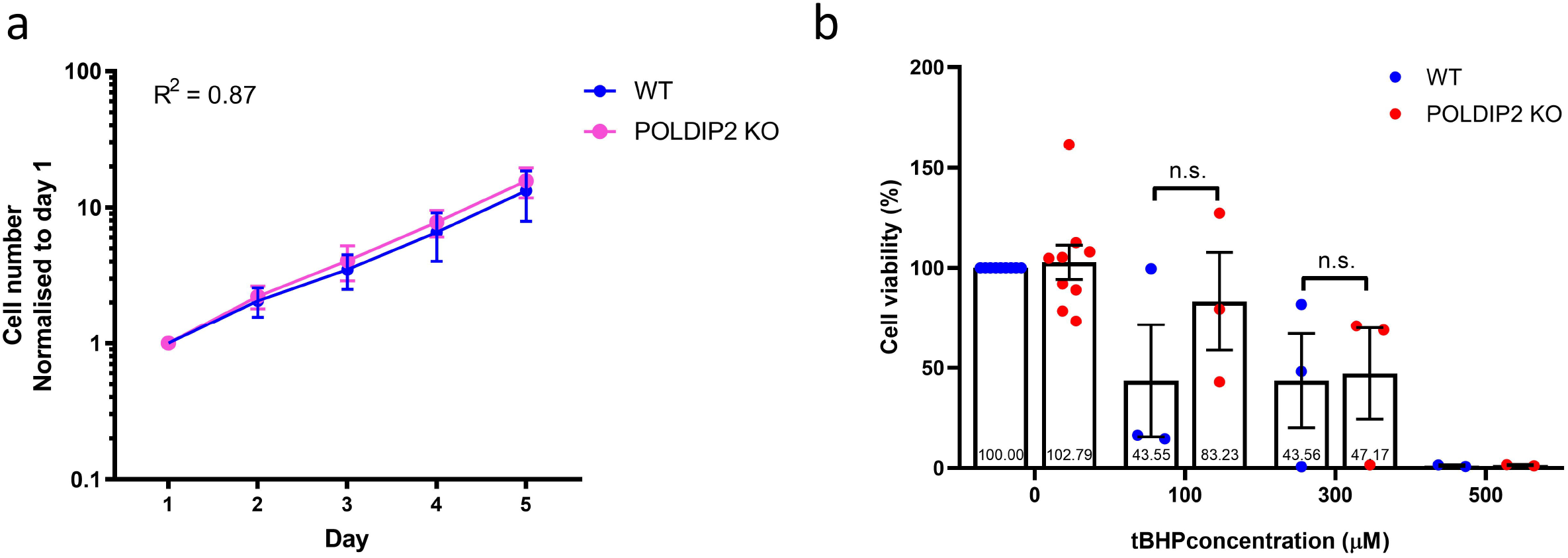
Knockout of *POLDIP2* in ARPE-19 shows normal levels of cell proliferation and viability. a) Cell proliferation of WT and POLDIP2 KO cell lines. Cell numbers expressed as mean ± SEM, n=3. b) Cell viability analysis of WT and POLDIP2 KO cell lines in the presence or absence of tBHP. Results are presented as mean ± SEM of 2-9 biological repeats, each with 8 technical repeats.

Oxidative stress plays an important role in AMD pathogenesis and progression ^29^. RPE has a high metabolic demand and thus mitochondria are a major source of ROS in the RPE. As a result, age-related mitochondrial dysfunction can induce oxidative stress in the RPE and thus leading to AMD ^30^. We assessed if *POLDIP2* knockout would affect cell viability of ARPE-19 in the presence of oxidative stress. To induce oxidative stress, we exposed the cells to tert-Butyl hydroperoxide (tBHP), a potent ROS-inducer commonly used to induce oxidative stress in cells and tissues. We then assessed cell viability of ARPE-19 treated with varying concentrations of tBHP. Our results showed that in the absence of tBHP, POLDIP2 KO cells showed a high level of cell viability and this level was comparable to WT (KO: 102.79±8.60% compared to WT, Figure 3b), which indicates that loss of *POLDIP2* did not affect cell viability. Following treatment with 100μM of tBHP, POLDIP2 KO cells exhibited higher cell viability compared to WT (83.23±24.42% and 43.55±28.04%, respectively), albeit this difference is not statistically significant. 300μM of tBHP caused similar levels of cell death in POLDIP2 KO and WT cells (47.17±22,76% and 43.56±23.51%, respectively), while 500μM of tBHP killed most of the cells in POLDIP2 KO and WT (1.48±0.31% and 1.23±0.40%, respectively). Overall, our results showed that *POLDIP2* knockout did not affect cell viability in the presence of oxidative stresses.

A previous study showed that *POLDIP2* knockout increased autophagy in mouse embryonic fibroblasts ^21^. Thus, we also investigated the role of *POLDIP2* in autophagy in ARPE-19 cells. We analysed LC3B levels as an indicator of autophagic flux ^31^ (Figure 4a). In the presence of lysosomal protease inhibitors Pepstatin and E64d controls, which partially inhibit degradation of LC3B-II, LC3B-II levels increased in both WT (4.22±0.83 compared to WT mock, Figure 4b) and POLDIP2 KO samples (2.70±0.82 compared to POLDIP2 KO mock: 1.40±0.31). Our results showed that LC3B-II levels in POLDIP2 KO cells were slightly higher than in WT (1.40±0.31 compared to WT, Figure 4b), albeit this difference is statistically insignificant. Similarly, LC3B-II/total LC3B ratio in POLDIP2 KO cells was slightly lower than in WT (0.80±0.07 compared to WT, Figure 4c), however this difference is also not statistically significant. Altogether, our results indicated that *POLDIP2* knockout did not significantly alter autophagic flux in ARPE-19.

**Figure 4.**
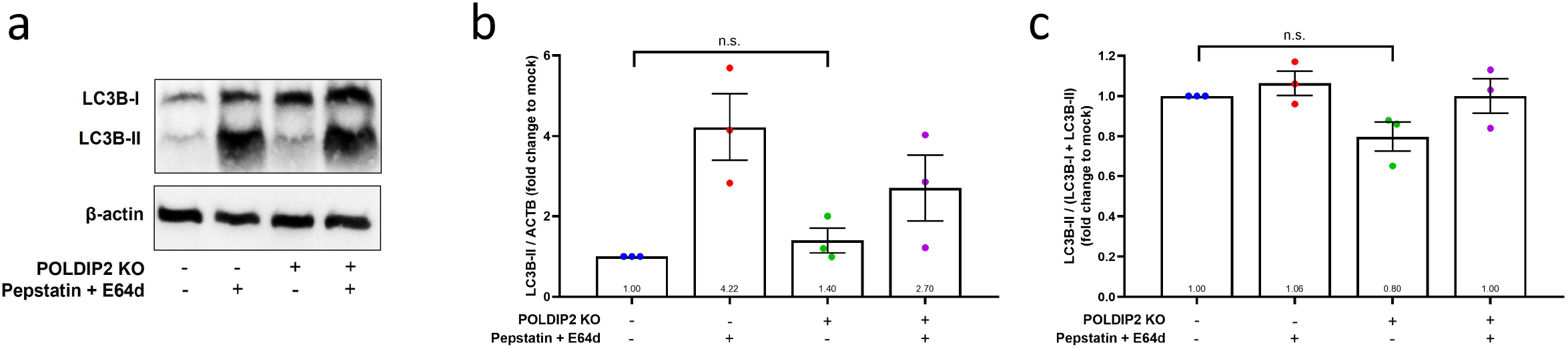
Knockout of POLDIP2 did not significantly alter autophagy in ARPE-19. a) Western blot analysis of LC3B protein levels in WT and POLDIP2 KO samples. E64d and pepstatin A protease inhibitors (10 mg/ml each) were added to the medium where indicated. β-actin serves as a loading control. b) Quantification of the ratio of LC3B-II to β-actin. Values expressed as mean ± SEM, n=3. n.s. not significant. c) Quantification of the ratio of LC3B-II/(LC3B-I + LC3B+II). Values expressed as mean ± SEM, n=3. n.s. not significant.

### Transcriptome profiling of WT versus POLDIP2 KO cell lines

To investigate the impact of *POLDIP2* knockout on RPE transcriptome profile, we performed RNA-seq on WT and POLDIP2 KO cell lines. Our results showed 93 upregulated genes and 203 downregulated genes in POLDIP2 KO compared to WT (Supplementary data). Figure 5a illustrated the top 50 differentially expressed (DE) genes between POLDIP2 KO and WT. We compared the expression levels of four RPE markers *BEST1, PMEL, RDH5*, and *RDH10* between the two cell lines (Figure 5b). Notably, the expression levels of RPE markers between POLDIP2 KO and WT were similar, indicating that the ARPE-19 retained RPE identity following the loss of *POLDIP2*.

**Figure 5.**
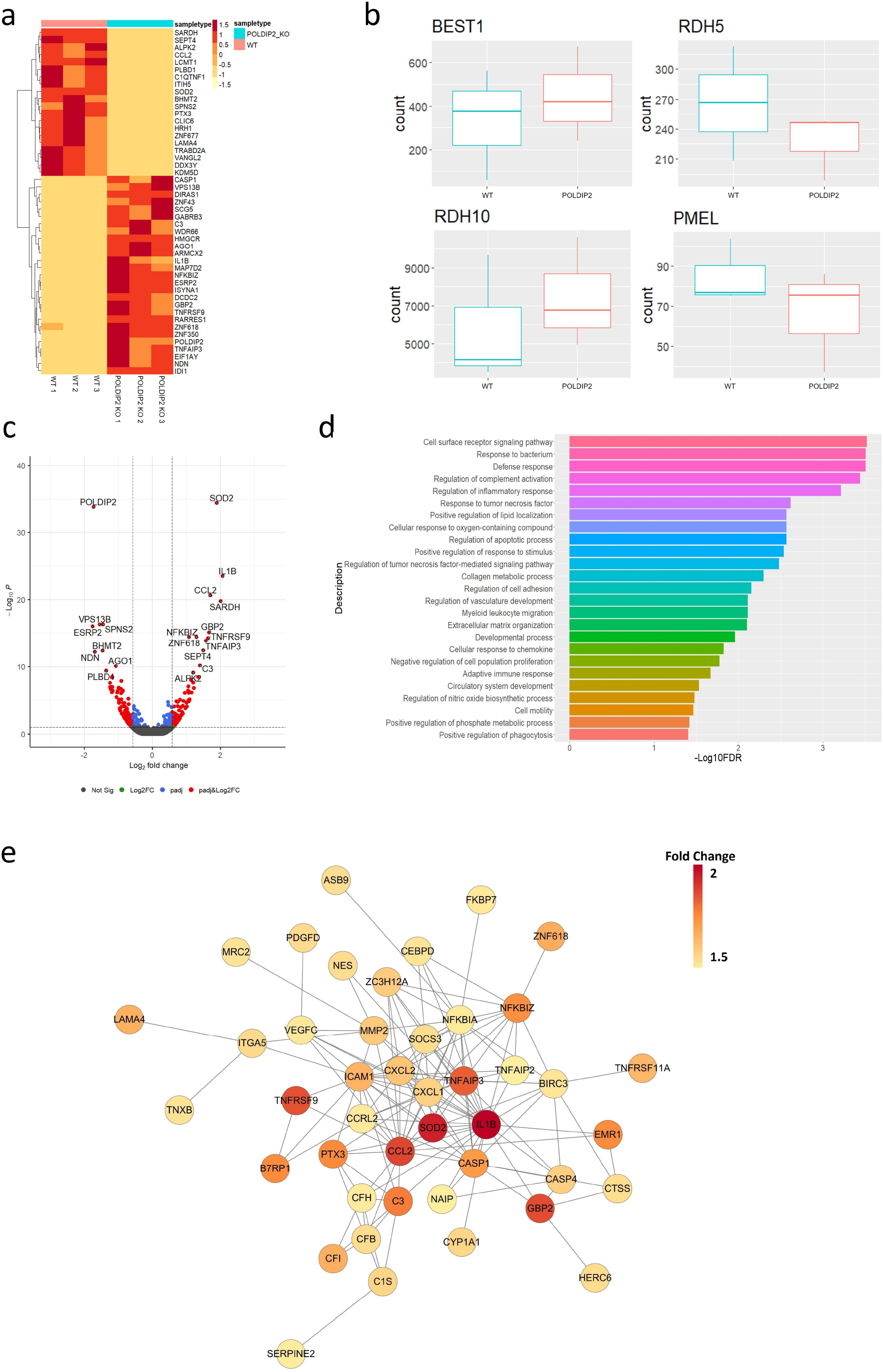
Transcriptome profiling of POLDIP2 knockout in ARPE-19. a) Heatmap of top 50 DE genes detected in WT (n=3) and POLDIP2 KO (n=3) cell lines. Changes in expression level shown from dark red to light yellow. b) Boxplot of the expression levels of RPE markers BEST1, RDH5, RDH10, and PMEL in different samples. c) Volcano plot of the top 20 DE genes labelled in POLDIP2 KO cell lines (n=3). d) GO annotation of the top 50 up-regulated DE genes in POLDIP2 KO samples (n=3). e) Network topology of the top up-regulated DE genes in POLDIP2 KO samples (n=3). Fold change value indicated from red to yellow.

As expected, we observed that the most down-regulated gene was *POLDIP2*, which confirms the quality of the knockout cell line (Figure 5c). Interestingly, the most upregulated gene was *SOD2* (Figure 5c). *SOD2* encodes manganese superoxide dismutase (MnSOD), an antioxidant enzyme located in the mitochondrial matrix that converts superoxide anion to hydrogen peroxide and protects mitochondria from oxidative stress ^32^. Knockdown of *SOD2* in RPE of mice induced oxidative damage, which led to morphological abnormalities in RPE and Bruch’s membrane, as well as other changes associated with AMD such as increase in autofluorescence levels and bis-retinoid pigments located in RPE and drusen, and accumulation of oxidatively modified proteins ^32^. Other top upregulated genes in POLDIP2 KO included *IL-1β, CCL2, SARDH, GBP2, NFKBIZ, TNFRSF9, TNFAIP3*, and *C3* (Figure 5c), most of which are involved in the immune defence system.

To elucidate the biological roles of the DE genes, we performed gene ontology analysis using the top 50 upregulated DE genes in POLDIP2 KO samples (Figure 5d). Interestingly, loss of *POLDIP2* also upregulated several genes involved in the complement system, including *C3, IL1B, CFI, CFH, CFB* and *C1S*. Critically, *C3, IL1B, CFI, CFH*, and *CFB* have been identified as genes implicated in AMD development ^11,33–36^. In addition, our results highlighted many upregulated DE genes are involved in the immune response, including chemokines (*CCL2, CXCL1, CXCL2*), cytokines (*IL1B*), and genes associated with cytokine-mediated signalling (*MMP2, ICAM1, GBP2, SOD2, NFKBIZ*), tumour necrosis factor-induced genes (*TNFRSF9, TNFAIP3*), and caspase cascade in apoptosis (*CASP1, CASP4*) (Figure 5e). Also genes involved in vasculature development and homeostasis were upregulated following *POLDIP2* loss (*LAMA4, VEGFC, SOCS3*, and *ZC3H12A*), as well as those involved in oxidative stress (*MMP2, ZC3H12A, SOD2*, and *TNFAIP3*). Furthermore, network topology analysis revealed the inter-connectivity between these DE genes, such as complement system genes (*CFB, CFH, C3, C1S, CFI*), and caspase cascade genes (*CASP1, CASP4*) (Figure 5e). Collectively, our results suggested that loss of *POLDIP2* affected genes involved in a wide range of biological processes, including various aspects of the immune response such as complement activation, and AMD-related processes such as vasculature development and oxidative damage.

### *POLDIP2* knockout reduced mitochondrial superoxide

Given the mitochondrial gene *SOD2* was the most-upregulated gene in POLDIP2 KO, we further studied the role of *POLDIP2* in regulating mitochondrial oxidative stresses and activity. We performed a MitoSox assay to compare the levels of mitochondrial superoxide between WT and POLDIP2 KO cell lines. In the presence of the positive control N-acetyl-L-cysteine (NAC), which inhibits oxidation, MitoSox fluorescence decreased in WT samples (0.70±0.04 compared to mock, Figure 6a). Interestingly, our analysis showed a significant reduction in mitochondrial superoxide levels in POLDIP2 KO (0.85±0.04 compared to WT, Figure 6a), which is consistent with the elevated expression of *SOD2* in POLDIP2 KO cells. In addition, we assessed the mitochondrial membrane potential using tetramethylrhodamine methyl ester (TMRM). Carbonyl cyanide 3-chlorophenylhydrazone (CCCP) was used as a positive control and reduced mitochondrial membrane potential in WT (0.65±0.06 compared to mock, Figure 6b). Importantly, we observed similar levels of mitochondrial transmembrane potential between WT and POLDIP2 KO cells (POLDIP2 KO: 1.07±0.03 compared to WT, Figure 6b). These results indicate that mitochondrial membrane potential was not affected by loss of *POLDIP2*. Altogether, our results identified a novel link of *POLDIP2* and *SOD2* in regulation of mitochondrial superoxide in RPE cells.

**Figure 6.**
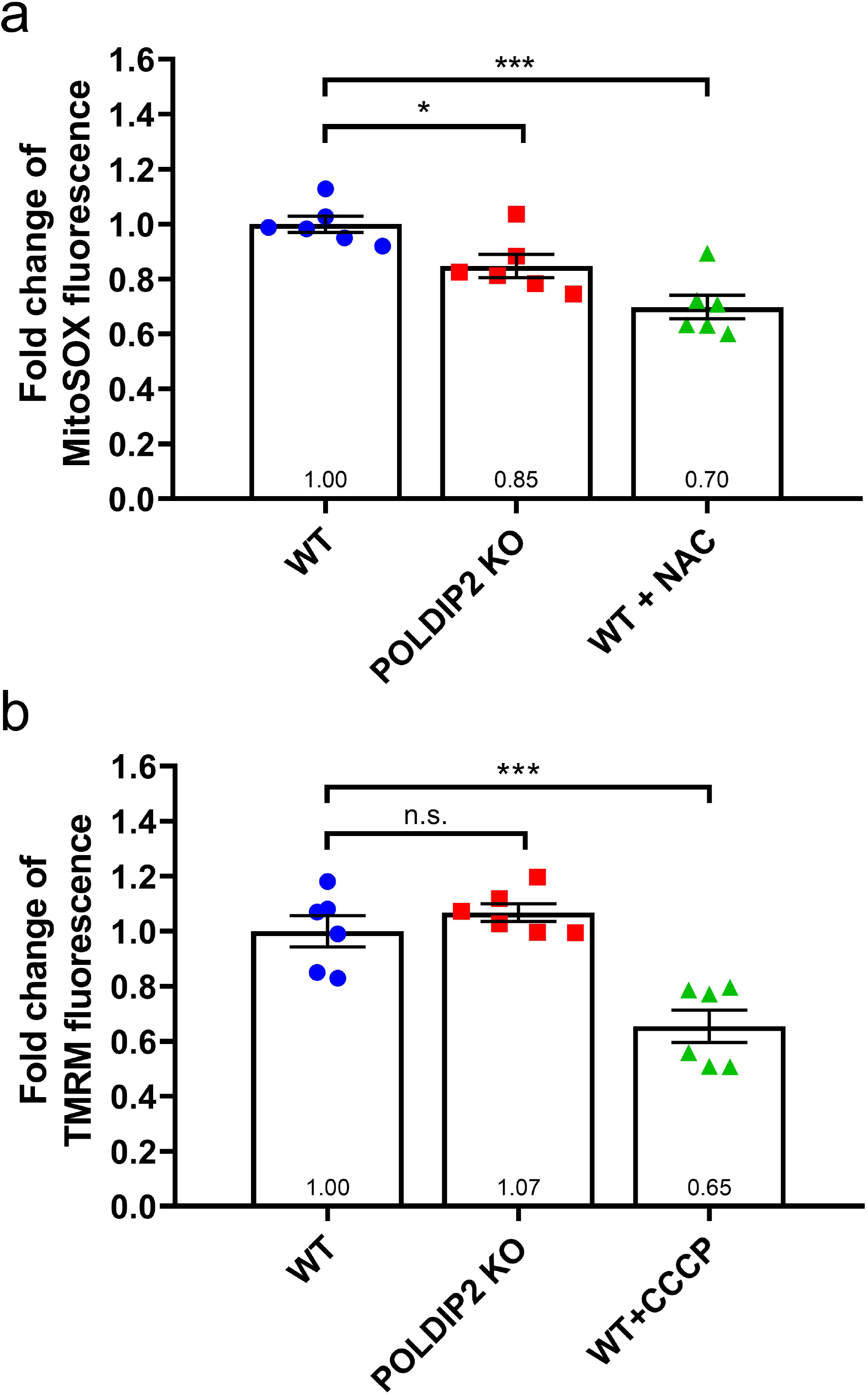
POLDIP2 KO reduced mitochondrial superoxide. a) Fold change of MitoSOX fluorescence in WT and POLDIP2 KO samples. N-acetyl-L-cysteine (NAC) was used as a positive control. b) Fold change of TMRM fluorescence in WT and POLDIP2 KO samples. Carbonyl cyanide 3-chlorophenylhydrazone (CCCP) was used as a positive control. Results are presented as mean ± SEM of 2 biological repeats, each with 3 technical repeats. n.s. not significant, * p<0.05 and *** p<0.001.

## Discussion

Considerable effort has been made to understand the genetic factors that contribute to AMD using retinal cell models. However, there are significant limitations in studying AMD in animal models. For example, many disease-associated signs only develop in aged rodents, which increases the length of time and subsequently the cost needed to house animals. Moreover, rats and mice do not have a macula and do not develop drusen or drusen-like deposits beneath the RPE ^37^. In this sense, simian primates are more appropriate animal models to study AMD, but are much more costly and require rigorous experiment setup ^38^. Consequently, *in vitro* models offer a cheaper and easier alternative to facilitate studies of gene function in the retina. ARPE-19 is commonly used to study retinal cell biology and shows similar features to native RPE cells, such as expression of transporters, barrier formation, and phagocytic ability ^39–41^. The present study reports the generation of an *in vitro* human RPE model with *POLDIP2* knockout using CRISPR as a model to study the function of an AMD-associated gene *POLDIP2*. Our first attempt to repress *POLDIP2* with a CRISPRi system yielded a low knockdown level, further optimisation of sgRNA would be important to improve this knockdown efficiency. We moved to a CRISPR KO system and generated a stable ARPE-19 cell line with *POLDIP2* knockout, providing an important tool to study the effect of *POLDIP2* on biological processes relevant to RPE cells and to AMD pathophysiology.

A previous study showed that Poldip2 affects growth rate and autophagy in MEFs ^21^. *Poldip2* knockdown caused markedly reduced growth in MEFs. Further investigations revealed that *Poldip2*-/-MEFs were arrested or delayed in both G1 and G2/M phases of the cycle, therefore decreasing the number of cells in S-phase and the protein levels of key cell cycle regulators also decreased following *Poldip2* knockdown. This study also reported increased autophagy following *Poldip2* knockout as indicated by higher levels of LC3B-II. However, we observed that loss of *POLDIP2* did not affect the growth rate and autophagic flux of ARPE-19 cells, suggesting that *POLDIP2* may have specific functions in different cell types.

To further study the roles of *POLDIP2*, we employed RNA-seq analysis to reveal gene expression changes caused by *POLDIP2* loss. We showed that genes related to the immune system were upregulated in the POLDIP2 KO group. *IL-1β* was the second most upregulated gene in POLDIP2 KO samples. *IL-1β* is transcribed following pro-inflammatory stimuli ^42^ and its expression is increased in the central nervous system in acute and chronic neurodegenerative diseases ^43^. In light-challenged mice, a model of dry AMD, subretinal mononuclear phagocytes expressed IL-1β, which induced rod death and cone segment loss ^34,44^. *CCL2* is another gene that was highly upregulated in POLDIP2 KO cells. CCL2 is a chemokine that directs leukocyte migration and its expression in RPE and the retina is very low in healthy young adult animals ^45^, but is elevated with ageing ^45^, following acute inflammation ^46,47^, and oxidative insult in the RPE ^48^. GBP2 is a member of the large GTPase superfamily that is strongly induced by interferon-γ (IFN-γ) and its expression in the retina was found to be significantly increased in aged and adult rats compared to young rats ^49^. Tumour necrosis factors *TNFRSF9* and *TNFAIP3* were also upregulated following *POLDIP2* knockout. These genes are associated with dendritic cell maturation, among which reduced levels of *TNFAIP3* was found to enhance dendritic cell function in patients with AMD ^50,51^. Our analysis also showed increased expression of the complement component *C3*, which is strongly associated with AMD ^33,52^ and has been shown to be interconnected with the expression of VEGF, RPE deterioration, geographic atrophy, and development of choroidal neovascularization ^53–55^.

Our findings revealed that following tBHP-induced oxidative stress, *POLDIP2* knockout did not lead to any remarkable changes to ARPE-19 viability. It has been reported that Poldip2 silencing in mouse embryonic fibroblasts increases cell sensitivity to oxidative stress as indicated by cell viability assay after H_2_O_2_ treatment^56^. While we observed normal cell viability in POLDIP2 KO cells following tBHP-induced oxidative stress in this present study, future studies could look at the effect of *POLDIP2* using other oxidative stress assays. Although loss of *POLDIP2* did not affect cell survival in the presence of external oxidative stress, we identified a link between *POLDIP2* KO and the reduced levels of mitochondrial superoxide. We hypothesise that loss of *POLDIP2* upregulated *SOD2*, which enhanced the conversion of superoxide anion to hydrogen peroxide and resulted in lower superoxide levels in the mitochondria. Previous studies have also demonstrated that *POLDIP2* is involved in oxidative signalling through cellular oxidases. Poldip2 has been reported to be an upstream regulator of the NADPH oxidase Nox4 ^57,58^, an enzyme functionally linked to proinflammatory responses, oxygen sensing, and senescence. In vascular smooth muscle cells, Poldip2 upregulates endogenous reactive oxygen species via Nox4 and positively regulates basal ROS production ^57^. In addition, Poldip2 also mediates oxidative stress and inflammation via interaction with Nox4 in lung epithelial cells and downregulation of Poldip2 leads to decreased production of ROS ^58^. It was demonstrated that Poldip2 deficiency protects against lung edema and vascular inflammation through suppressing mitochondrial ROS in a mouse model of acute respiratory distress syndrome ^59^. Furthermore, deletion of *Sod2* in mice has been shown to disrupt RPE morphology, reduce RPE function, and elevate oxidative stress in RPE ^6^. This study supports the notion that mitochondrial damage plays a role in RPE dysfunction and progression of AMD.

There are limitations to this study. Even though ARPE-19 has morphological and functional features of human RPE, they have their own limitations. The loss of RPE melanin has been reported in AMD ^60^, and since ARPE-19 cells do not retain the original pigmented phenotype of the RPE, this hinders an opportunity to study changes in melanin pigment in RPE. Future studies using primary RPE cells or stem cells-derived RPE would provide a suitable model to validate the functions of *POLDIP2* in RPE cells.

In summary, we have generated a *POLDIP2* knockout ARPE-19 cell line using CRISPR/Cas9 and studied the biological functions of *POLDIP2*. To our knowledge, this is the first functional study of *POLDIP2* in retinal cells to understand its potential role in AMD. The POLDIP2 KO cell line possesses normal proliferation, phagocytosis, autophagy, sensitivity to oxidative stress-induced cell death and mitochondrial activity. Interestingly, we identified a novel link between *POLDIP2* and mitochondrial oxidative stress modulation via *SOD2*, supporting a potential role for *POLDIP2* in AMD pathogenesis. Future studies to investigate the precise mechanism by which *POLDIP2* regulates oxidative stress signalling pathways would be important to advance our understanding of AMD genetics.

## Methods

### Cell culture

HEK293FT, ARPE-19, ARPE-19-KRAB, and POLDIP2 KO cells were maintained at 37°C and 5% CO2 in a culture medium containing DMEM (Thermo Fisher) supplemented with 10% [v/v] Fetal Bovine Serum (FBS), 2mM GlutaMAX, and 0.5% Penicillin-Streptomycin (all from Thermo Fisher). Cells were passaged before they reached confluency using 0.25% Trypsin-EDTA (Thermo Fisher).

### Generation of POLDIP2 knockout ARPE-19 cell line

For construction of a lentiviral vector co-expressing SpCas9 and target sgRNA, the complementary DNA oligos of the sgRNAs targeting *POLDIP2* were commercially synthesised, then phosphorylated and annealed using T4 Ligation Buffer and T4 PNK (NEB) to form the double-strand DNA using the following thermal parameters: at 37°C for 30 min, and at 95°C for 5 min, followed by decreasing at 5°C/min to 25°C. The lentiviral vector lentiCRISPRv2 (Addgene, #52961) was linearised by BsmBI and dephosphorylated using CIP (NEB). The product was purified by gel electrophoresis and gel extract. The annealed DNA oligos of the sgRNA was ligated to the linearised lentiCRISPRv2 vector by T4 DNA ligase (NEB).

For lentivirus generation, 7×10^6^ HEK293FT cells were seeded in a 10 cm^2^ dish one day prior to transfection, cultured in Opti-MEM supplemented with 5% FBS and 200 μM Sodium pyruvate (all from Thermo Fisher). Lentivirus was generated using a 3^rd^ generation packaging system. The transfer plasmid and three packaging vectors pMDLg/pRRE (Addgene, #12251), pRSV-Rev (Addgene, #12253), and pMD2.G (Addgene, #12259) were transfected into HEK293FT cells using Lipofectamine 3000 (Thermo Fisher). 6 hours after transfection, the medium containing Lipofectamine 3000 was replaced with fresh media. The supernatant containing the virus was collected 48 and 72 hours after transfection, and subsequently filtered (0.45 μm filter, Sartorius) and concentrated using PEG-it precipitation solution (SBI Integrated Sciences) according to the manufacturer’s instructions. Virus was resuspended in cold PBS and titre was determined using Lenti-X p24 Rapid Titre Kit (Takara Bio) according to manufacturer’s instructions.

ARPE-19 cells were transduced with the POLDIP2 lentiviruses (MOI=10) overnight, followed by selection with 2μg/ml puromycin 3 days after transduction. Transduced ARPE-19 cells were further expanded to obtain the POLDIP2 KO cell line.

### RT-qPCR analysis

RNA extraction was performed using illustra RNAspin Mini Kit (GE Healthcare Life Sciences) according to manufacturer’s instructions. RNA concentration and quality were measured using NanoDrop. cDNA was synthesised using High-capacity cDNA reverse transcription kit with RNase inhibitor (Thermo Fisher). RT-qPCR was reaction mixture was set up using TaqMan Fast Advanced Master Mix (Thermo Fisher) and Taqman probes for POLDIP2 (Hs00210257_m1) and the housekeeping gene β-actin (Hs99999903_m1) (Thermo Fisher). RT-qPCR was performed on the 7500 Fast or StepOnePlus Real-Time PCR System (Thermo Fisher), following manufacturer’s instructions. The delta delta Ct method was used to calculate and compare relative mRNA levels to control.

### Cell proliferation assay

On day 0, 1.2×10^4^ ARPE-19 WT and POLDIP2 KO cells were seeded in a well of a 24-well plate. Cells from three wells per cell line were harvested daily for the next five days using 0.25% Trypsin-EDTA, stained with trypan blue (Thermo Fisher), and counted using a Countess Automated Cell Counter (Thermo Fisher). The average number of live cells was calculated for each day and normalised to day 1 cell number.

### Cell viability assay

Cell viability assay was performed using a CellTiter-Glo Luminescent Cell Viability Assay (Promega) following the manufacturer’s instructions. On day 0, 10^4^ ARPE-19 and POLDIP2 KO cells were seeded in a well of a 96-well plate. On day 2, the cells were treated with various concentrations of tert-Butyl Hydroperoxide (tBHP) (Sigma). On Day 3, old media was replaced with 25μl of fresh media and 25μl of CellTiter-Glo Reagent was added to each well. The plate was incubated at room temperature for 10 minutes to generate a luminescent signal. 40μl of cell lysate from each well was loaded to an opaque white luminometer plate and luminescence was recorded using a Spark 20M microplate reader (Tecan). The OD reading was normalised to the control condition (without tBHP and gene KO) and results are presented as fold change relative to mock.

### Immunoblotting

Protein levels were assessed by Western blotting. Cells were lysed using RIPA Buffer (Thermo Fisher) and sonicated. Protein concentrations were determined using Pierce BCA Protein Assay Kit (Thermo Fisher). Protein lysates were mixed with 4X Laemmli Sample Buffer (Biorad, #1610747) and 2-Mercaptoethanol (1:40, Sigma) and heated at 95°C for 5 minutes. Proteins were separated via 15% SDS-PAGE gels and transferred to PVDF membranes. Following blocking, blots were incubated overnight with primary antibodies: anti-beta actin (1:2500 in 5% BSA, Abcam), anti-LC3B (1:1000 in 5% BSA, Cell Signaling), and anti-POLDIP2 (1:1000 in 5% BSA, Abcam). Membranes were then incubated with secondary antibodies (diluted 1:2500 in nonfat dry milk): goat anti-rabbit IgG HRP or goat anti-mouse IgG HRP (all from Abcam). Bands were visualised using Pierce ECL Western Blotting Substrate (Thermo Fisher) in a BioRad Chemidoc MP Imaging System. Band quantification was performed using ImageJ.

### Phagocytosis

4×10^5^ cells were seeded in a well of a 6-well plate. Cells were incubated with 1μm diameter, yellow-green (505/515 nm) carboxylate-modified microspheres (FluoSpheres, Thermo Fisher) at a quantity of 160 beads per cell for 4 hours. Cells were dissociated with 0.25% Trypsin-EDTA, washed with DPBS + 1% FBS 5 times, and resuspended in 400μl of DPBS + 1% FBS. The cell samples were added with 0.1 μg/ml of DAPI (Sigma) and passed through a strainer (In Vitro Technologies). Quantification of FITC+ cells was performed using a BD LSRFortessa Cell Analyzer (BD Biosciences). Gating was set with a negative control using WT cells without FluoSpheres.

### Mitochondrial membrane potential assay

Mitochondrial membrane potential was assessed using tetra-methyl rhodamine methyl ester (TMRM), which selectively accumulates in the mitochondria according to the mitochondrial membrane potential. Cells were incubated with a non-quenching dose of TMRM at 10 nM in culture media. The mitochondrial respiratory uncoupler Carbonyl cyanide 3-chlorophenylhydrazone (CCCP, 50 mM), was used as a positive control to dissipate the mitochondrial membrane potential. Images were captured at 200× magnification with a fluorescence microscope (Olympus IX71) and the total corrected cell fluorescence was assessed using ImageJ. At least 600 cells from 3 random fields were counted per group.

### Mitochondrial superoxide production assay

Mitochondrial production of reactive oxygen species (ROS) was assessed using MitoSOX Red (Thermo Fisher Scientific). Cells treated with 5 mM of the antioxidant N-acetyl-L-cysteine (Sigma-Aldrich) were used as a positive control. Images were captured at 200× magnification with a fluorescence microscope (Olympus IX71) and the total corrected cell fluorescence was assessed using ImageJ. At least 600 cells from 3 random fields were counted per group.

### RNA sequencing

Total RNA of ARPE-19 cell lines was extracted using the Illustra RNAspin Mini Kit (GE Healthcare Life Sciences) according to manufacturer’s instructions. RNA quality was checked by bioanalyzer and the TruSeq Stranded mRNA kit (Illumina) was used to prepare transcriptome libraries. The libraries were sequenced using Illumina Novaseq 6000 100bp single-end sequencing, at a depth of 38-50 million reads per sample (Australian Genome Research Facility).

Following the abundance estimates of transcripts generated by *Salmon* v1.8, the pseudocounts were mapped to the GRCh38 genome assembly using the *tximport* v1.22.0 package ^61^. The gene count matrix was inputted as an DESeq2Dataset object using the *DESeqDataSetFromTximport* function, then the DESeq2Dataset object was normalised using the *counts* function to make fair gene expression comparisons between samples ^62^. The normalised dataset was analysed with the *DESeq2* v1.34.0 R package using rlog transformation. The sample-level QC was performed using principal components analysis while the gene-level QC was performed using hierarchical clustering. For differential expression analysis, the significant differentially expressed genes were determined using the *filter* function with adjusted p value of < 0.05 and fold change > 1.5. The expression data of significant differentially expressed genes was visualised using the *ggplot2* v3.3.6, *pheatmap* v1.0.12 and *EnhancedVolcano* v1.12.0 R package ^63–65^. The upregulated DE genes were used for network topology analysis using Cytoscape v3.8 ^66^, with default setting of full STRING network, a confidence score cutoff of 0.4 and no additional interactor, resulting in a network with 44 DE genes. Gene ontology analysis was performed for the upregulated DE genes using functional enrichment analysis in Cytoscape with default settings.

### Short tandem repeat analysis

Genomic DNA of ARPE-19 cell lines was extracted using the Wizard SV Genomic DNA Purification System (Promega), following manufacturer’s instructions. Short tandem repeat analysis was performed using the GenePrint 10 system (Promega) by the Australian Genome Research Facility.

### Statistical analysis

Data are presented as the mean ± standard error of the mean (SEM) of at least three independent experiments. p<0.05 is used to establish statistical significance. RT-qPCR for *POLDIP2* expression, phagocytosis, cell viability, and autophagy assays were assessed using unpaired t-test, mitochondrial assays were analysed using one-way ANOVA (GraphPad Prism).

## Data availability

The transcriptome data generated in this study are available in the NCBI Gene Expression Omnibus database (GSE207158), including raw data, processed data, information of the experimental design, sequencing and processing pipeline.

## Author contribution

Conceptual design: TN, CL, RG, SYL, SSCH, AWH and RCBW; Conduct experiments: TN, DU, LW, JGL, JW, SSCH; Data analysis: TN, DUC, LW, JGL, SSCH, SYL, CL, RG, RCBW; Funding: CL, RG, TE, SYL, RCBW; Manuscript writing: TN, CL, RG, RCBW. All authors approved the manuscript.

## Acknowledgement

This work was funded by the University of Melbourne (RCBW), the Centre for Eye Research Australia (RCBW), and the Stafford Fox Medical Research Foundation (SYL). TN and DU are supported by the Melbourne Research Scholarship from the University of Melbourne. RG is supported by the National Health and Medical Research Council Fellowship. The Centre for Eye Research Australia and St Vincent’s Institute of Medical Research receive operational infrastructure support from the Victorian Government.

