## Supplementary fig 1-4 and supplementary table 1 for "Knockout of AMD-associated gene *POLDIP2* reduces mitochondrial superoxide in human retinal pigment epithelial cells"

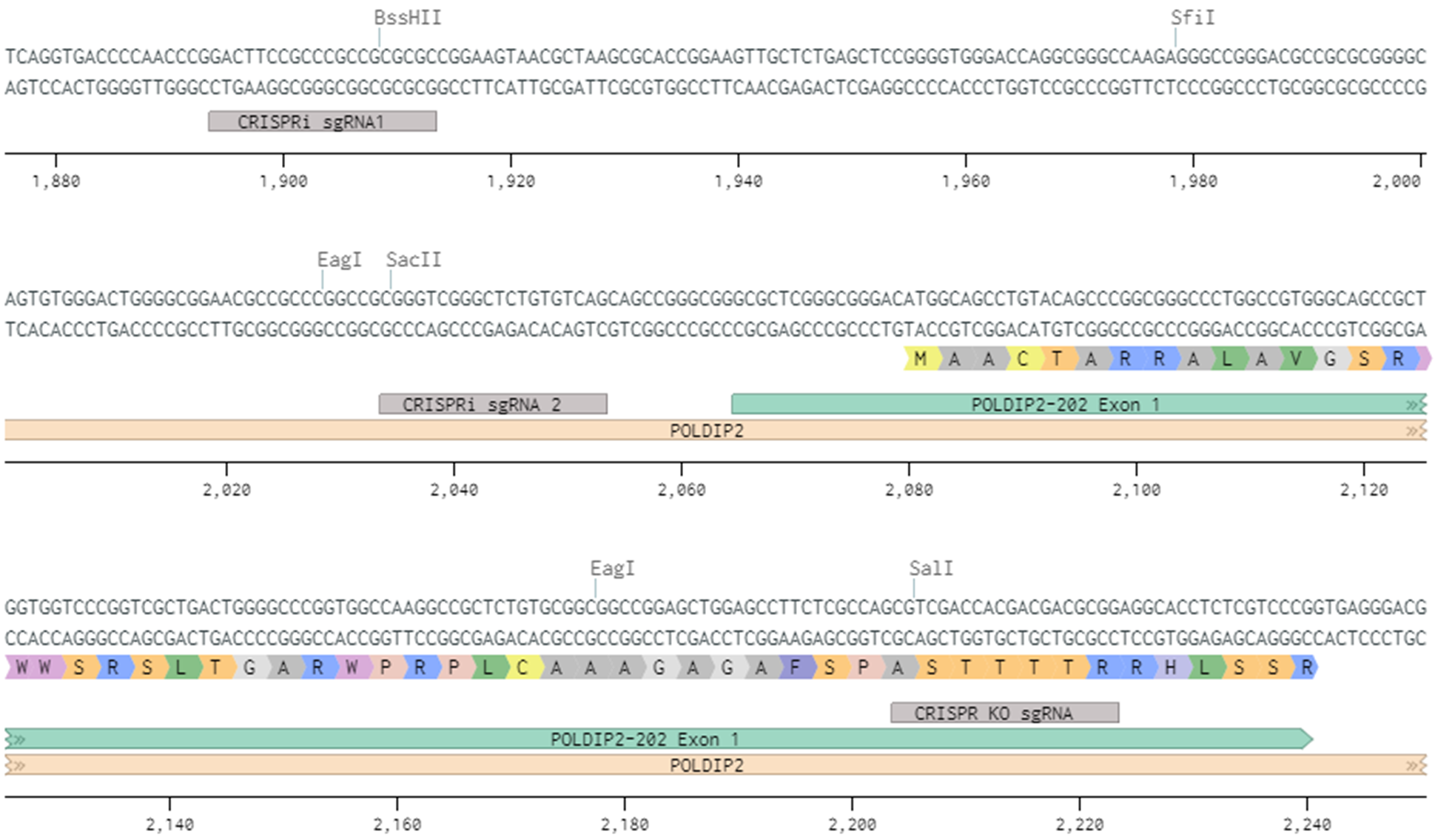

**Supplementary figure 1.** Schematic diagram of the 5’ region of human *POLDIP2* gene with sgRNA target areas near the transcription start site (TSS). CRISPRi sgRNA1 and CRISPRi sgRNA2 were used for *POLDIP2* knockdown and CRISPR KO sgRNA was used for *POLDIP2* knockout.

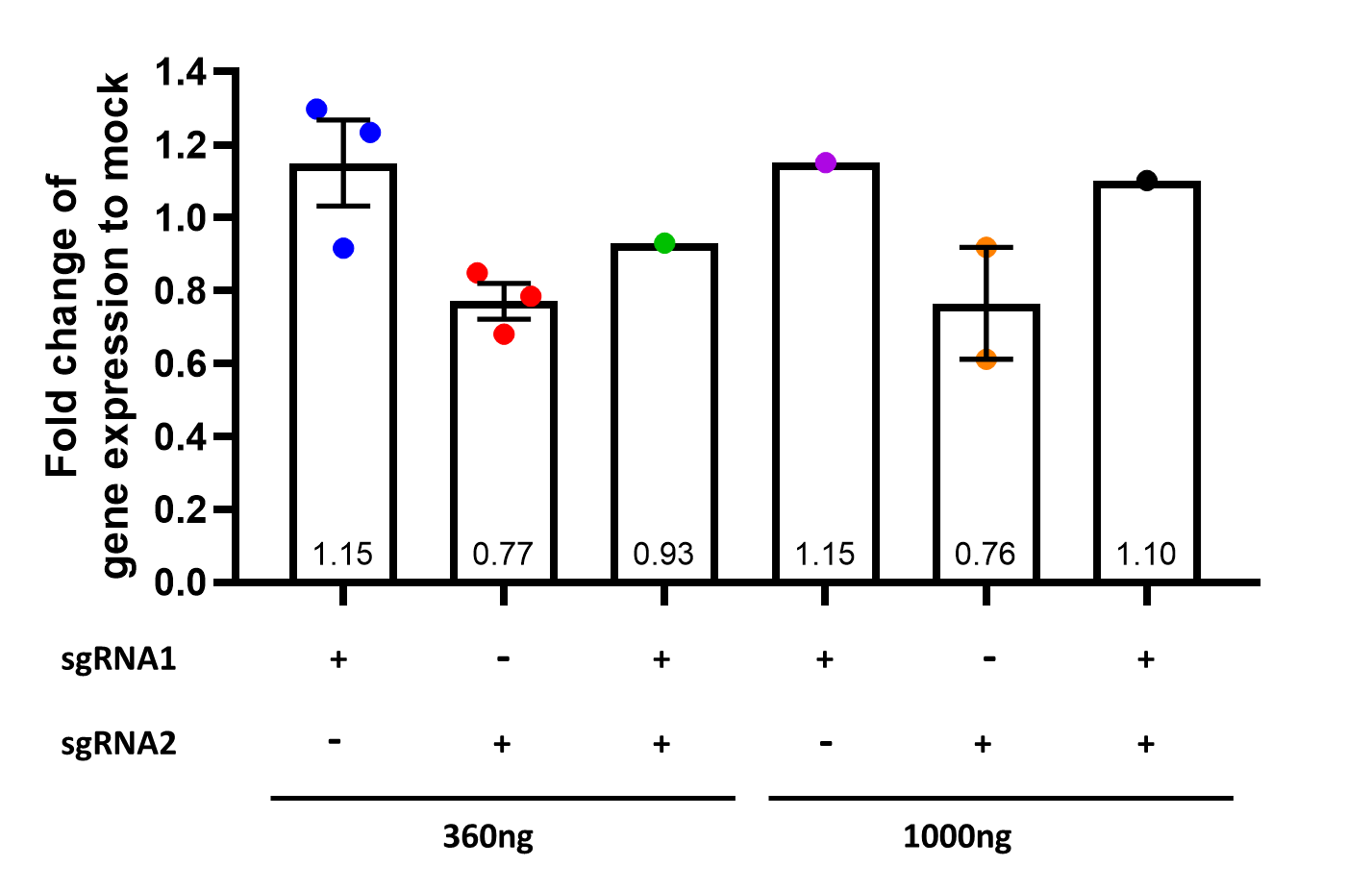

**Supplementary figure 2.** *POLDIP2* repression in ARPE-19-KRAB cell line. Cells were analysed using RT-qPCR 3 days after transfection with the indicated sgRNAs. Values were normalised to a wild type ARPE-19 mock control and expressed as mean ± SEM.

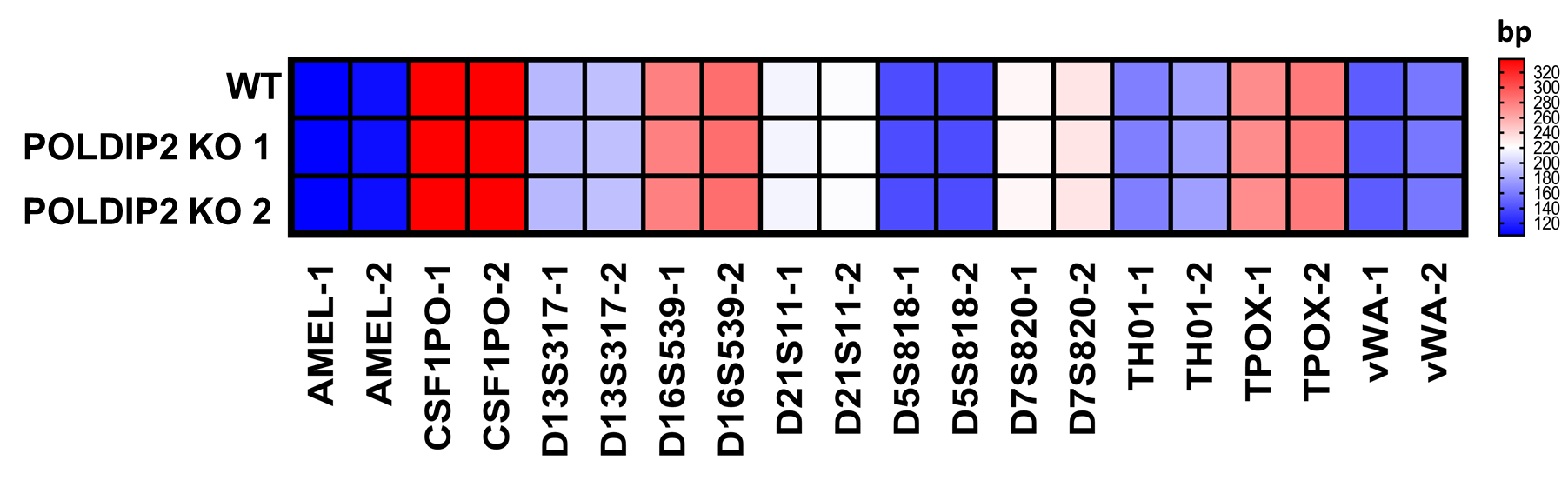

**Supplementary figure 3.** Heatmap of short tandem repeat analysis of 10 polymorphic markers of WT ^28^ and POLDIP2 KO cell lines. Allele 1 and 2 are designated as -1 and -2 respectively.
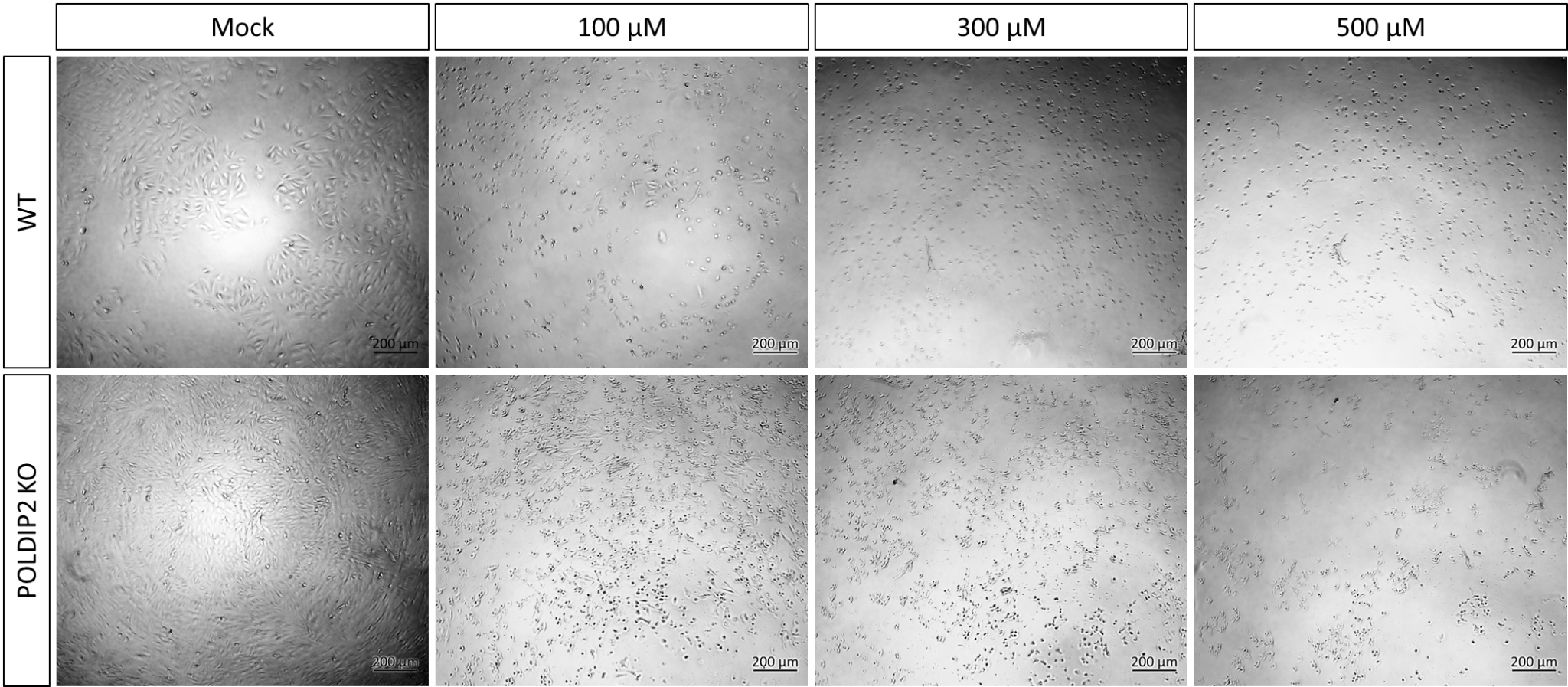

**Supplementary figure 4.** Representative images of WT and POLDIP2 KO cells after 3 days of tBHP treatment with various concentrations (100 μM, 300 μM, and 500 μM) and control (Mock).

**Supplementary table 1:** Information of sgRNAs used in this study. TSS distance is based on the transcription start site defined by Ensembl. On-target and off-target scores are based on ^67^.

| **Name** | **TSS distance** | **Strand** | **Sequence** | **PAM** | **On-target score** | **Off-target score** |
| --- | --- | --- | --- | --- | --- | --- |
| POLDIP2 CRISPRi sgRNA1 | 151 | - | GACTTCCGCCCGCCGCGCGC | CGG | 34.5 | 83.4 |
| POLDIP2 CRISPRi sgRNA 2 | 11 | - | CTGACACAGAGCCCGACCCG | CGG | 64.0 | 49.0 |
| POLDIP2 CRISPR KO | 139 | + | CGTCGACCACGACGACGCGG | AGG | 68.2 | 93.1 |
